# Classification-based Inference of Dynamical Models of Gene Regulatory Networks

**DOI:** 10.1101/673137

**Authors:** David A. Fehr, Manu, Yen Lee Loh

## Abstract

Cell-fate decisions during development are controlled by densely interconnected gene regulatory networks (GRNs) consisting of many genes. Inferring and predictively modeling these GRNs is crucial for understanding development and other physiological processes. Gene circuits, coupled differential equations that represent gene product synthesis with a switch-like function, provide a biologically realistic framework for modeling the time evolution of gene expression. However, their use has been limited to smaller networks due to the computational expense of inferring model parameters from gene expression data using global non-linear optimization. Here we show that the switch-like nature of gene regulation can be exploited to break the gene circuit inference problem into two simpler optimization problems that are amenable to computationally efficient supervised learning techniques. We present FIGR (Fast Inference of Gene Regulation), a novel classification-based inference approach to determining gene circuit parameters. We demonstrate FIGR’s effectiveness on synthetic data as well as experimental data from the gap gene system of *Drosophila*. FIGR is faster than global non-linear optimization by nearly three orders of magnitude and its computational complexity scales much better with GRN size. On a practical level, FIGR can accurately infer the biologically realistic gap gene network in under a minute on desktop-class hardware instead of requiring hours of parallel computing. We anticipate that FIGR would enable the inference of much larger biologically realistic GRNs than was possible before. FIGR Source code is freely available at http://github.com/mlekkha/FIGR.

## 1 Introduction

Development is controlled by gene regulatory networks (GRNs) that interpret extrinsic signals to make decisions about cell fate [6, 23]. The same principle also applies to the maintenance of populations of replenishable cell-types such as blood cells during adult homeostasis or to the *in vitro* reprogramming of cell identity [11]. Transcription factors (TFs) provide the connectivity in GRNs since they modulate the transcription rate of target genes by binding specific sites in non-coding DNA neighboring the gene and recruiting cofactor proteins that interact with the polymerase holoenzyme complex or modify the accessibility of DNA. Unlike direct regulation by TFs, non-TF gene products can regulate target gene transcription indirectly. For example, the abundance of receptor proteins modulates the transduction of extracellular signals [30].

Inferring and modeling the genetic control logic of cell-fate determination is necessary not only for understanding the basic principles of development and physiology but also for further progress in regenerative medicine. Over the past decade or so, it has become clear that developmental GRNs comprise tens to hundreds of densely interconnected genes [7, 29] rather than a few so-called master regulators. Moreover, developmental GRNs are wired recursively since the genes encoding TFs are themselves regulated by other TFs or indirectly by non-TF genes products. Their large size and high interconnectivity make the inference and predictive modeling of developmental GRNs a challenging problem.

GRNs can be reverse engineered from gene expression data sampled in time, in different cell types, or in different mutant backgrounds with a variety of statistical approaches [20, 26, 33]. One specific reverse-engineering methodology, termed *gene circuits*, has been particularly successful in inferring and modeling developmental GRNs [15, 31, 38]. Gene circuits model protein or mRNA concentrations using coupled nonlinear ordinary differential equations (ODEs) in which synthesis is represented as a switch-like function of regulator concentrations. The values of the free parameters define GRN connectivity. Gene circuits do not presuppose any particular connectivity, but instead determine it by estimating the values of the parameters from data using global nonlinear optimization techniques [5, 21, 22]. In contrast to other reverse engineering methods, gene circuits infer not only the topology of the network but also the type, either activation or repression, and strength of interactions. Most importantly, the inference procedure yields ODE models that can be used to simulate and predict developmental perturbations [16, 24, 25, 37, 39].

Despite its successes, the gene circuit method suffers from two drawbacks that have hamstrung its application to larger networks and other developmental systems. The first limitation is that gene circuit parameter inference is computationally intensive and scales with *G*^3^, where *G* is the number of genes. Because of this, gene circuits have only been inferred for relatively small networks of fewer than 10 genes so far [4, 24, 27]. The second limitation is that, parameter identifiability is relatively poor and only the sign of interaction, but not its strength, is inferable for the majority of network connections [2] in apparently well-determined problems.

In this paper we present an alternative approach, FIGR (Fast Inference of Gene Regulation), for determining gene circuit parameters that overcomes the first drawback by being significantly more computationally efficient than simulated annealing. Our algorithm exploits the observation that the inference of the connectivity of a given gene can be rephrased as a supervised learning problem: to find a hyperplane in phase space that classifies observations into two groups, one where the gene is ON and the other where the gene is OFF. Our algorithm determines whether a gene is ON or OFF at a given observation point by computing the time derivative of concentrations in a numerically robust manner. It then performs classification using either logistic regression or support vector machines (SVM) to determine the equation of the switching hyperplane. The genetic interconnectivity can then be computed from the coefficients of the hyperplane equation in a straightforward manner. We have implemented the algorithm in MATLAB and tested its ability to recover the genetic interconnectivity of random GRNs from simulated data. The algorithm works as expected and recovers parameters accurately. We also demonstrate the ability of our method to correctly infer the *Drosophila* gap gene regulatory network from empirical data. We observed a ∼600-fold speed up relative to simulated annealing on the gap gene problem. Finally, we discuss how conceiving of and visualizing gene circuit inference as a classification problem promises to provide insight into parameter identifiability.

## 2 Materials and Methods

### 2.1 Gene circuit models of GRNs

Gene circuits [31] describe the time evolution of the concentrations of *G* gene products *x*_*g*_(*t*) according to *G* coupled ordinary differential equations,

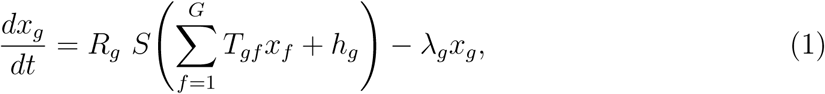

where *S*(*u*) is a sigmoid regulation-expression function,

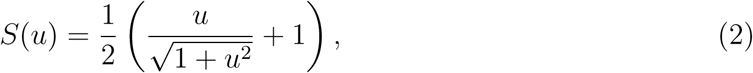

representing the switch-like nature of gene regulation. *R*_*g*_ is the maximum synthesis rate of product *g, T*_*gf*_ are genetic interconnectivity coefficients describing the regulation of gene *g* by the product of gene *f, h*_*g*_ determines the basal synthesis rate, and *λ*_*g*_ is the degradation rate of product *g*. Nominally, all genes in the model also function as regulators, so that both *g* and *f* run over the range 1, 2, 3, …, *G*. Sometimes such gene networks include upstream regulators that are not themselves influenced by other gene products represented in the model. For example, in the *Drosophila* segmentation gene network, maternal proteins such as Bicoid activate the zygotically expressed genes, but are not regulated by their targets [1]. An upstream regulator *g* can be represented by setting *T*_*gf*_ = 0 for all *f*.

#### 2.1.1 Glass networks

Here, we utilize gene circuits in which gene state switching is represented with the Heaviside regulation-expression function

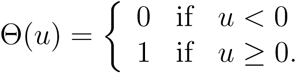

The resulting differential equations are piece-wise linear and are referred to as Glass networks [8, 9]. Glass networks are capable of exhibiting a rich variety of dynamical behaviors such as multistability, limit cycles, and chaos [10, 28].

Using the state vector **x** = (*x*_1_, *x*_2_, …, *x*_*G*_) to represent a point in the *G*-dimensional phase space of the model and the vector **T**_*g*_ to represent the *g*th row of the genetic interconnectivity matrix, the Glass equations may be written as

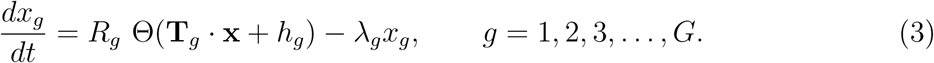

The gene may said to be “ON” or “OFF” depending on whether the gene product is being synthesized or not respectively. Equation (3) implies that

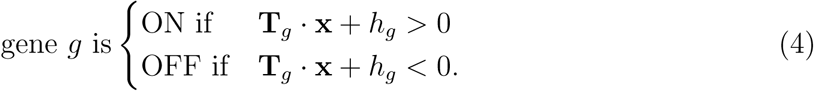

Thus the “gene *g* ON” and “gene *g* OFF” configurations are separated in phase space by the hyperplane defined by the equation **T**_*g*_ · **x** + *h*_*g*_ = 0. We call this the *switching hyperplane*. **T**_*g*_ is the normal to the switching hyperplane and **T**_*g*_ · **x** + *h*_*g*_ is the perpendicular distance of any point **x** to the hyperplane. Furthermore,

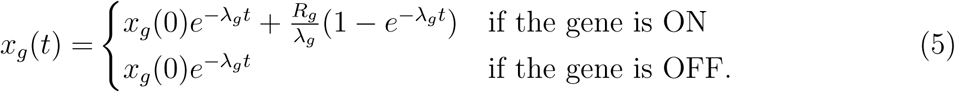

Equations (4) and (5) imply that for Glass networks, the *regulatory parameters*, **T** and *h*_*g*_, and the *kinetic parameters, R*_*g*_ and *λ*_*g*_, are separable. The former determine the switching hyperplane, while the latter determine the trajectories on either side of the hyperplane. Figure 1D,G shows examples of the switching hyperplanes and trajectory of a two-gene gene circuit (Fig. 1A,B) having a stable spiral equilibrium solution.

**Figure 1:**
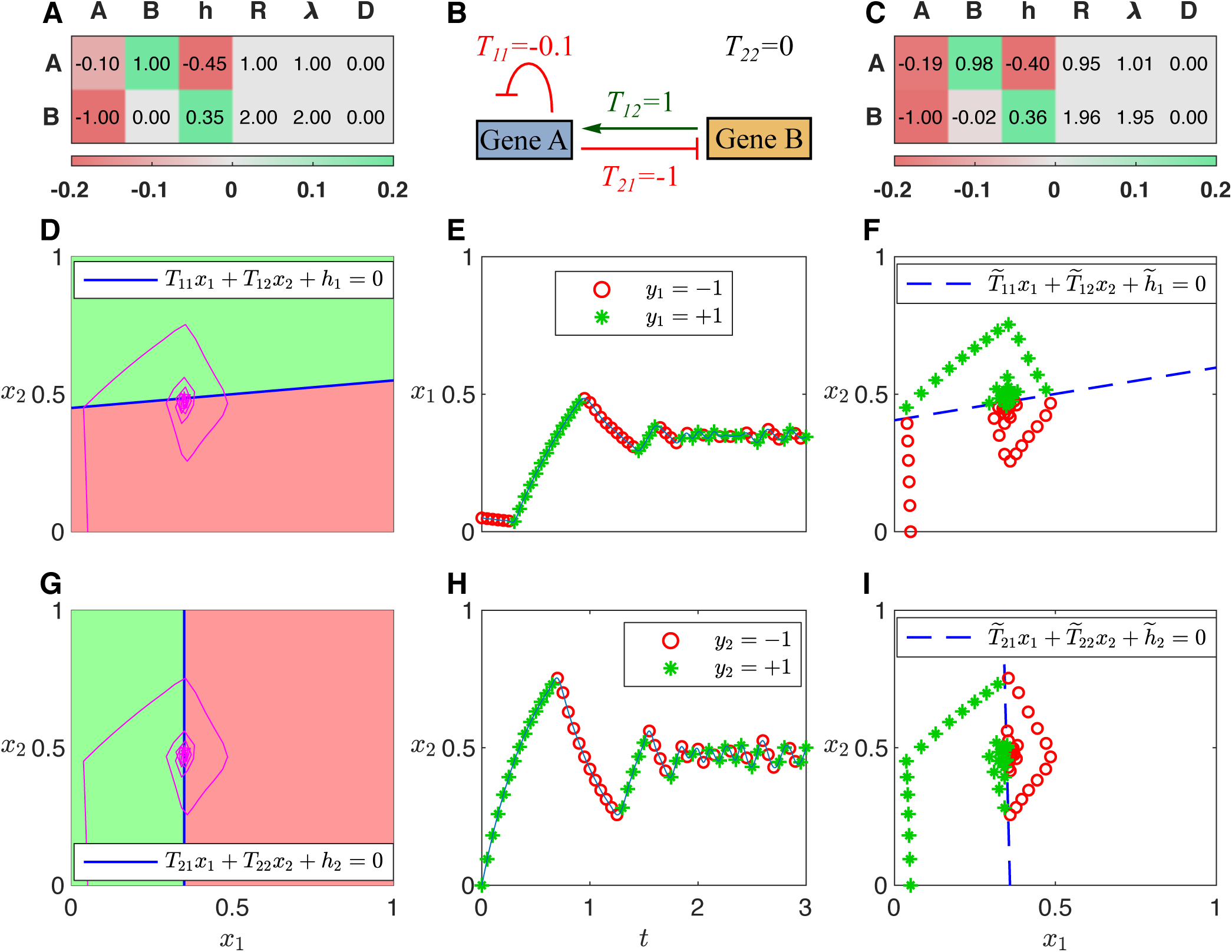
Classification-based inference of an example gene circuit. **A**. Theoretical parameter values are listed by row for each gene. The **T** matrix is shown in the first two columns, one column per regulator. Green (red) indicates activation (repression). **B**. Schematic of the theoretical gene circuit. **C**. Parameters inferred by FIGR. **D**,**G**. Trajectory in phase space (purple) overlaid upon the Heaviside regulation-expression function. Green (red) is ON (OFF). The switching hyperplane is plotted as a blue line. Switching hyperplanes for genes A and B are showing in panels D and G respectively. **E**,**H**. Sampled gene expression trajectories and assignment of ON/OFF state for genes A (panel E) and B (panel H). Trajectories are numerical solutions of Equation (3). Detected ON or OFF state (Section 2.2.1) is indicated with green stars or red circles respectively. **F**,**I**. Switching hyperplane (dashed blue line) inferred using Logistic regression for genes A (panel F) and B (panel I). Sampled trajectories annotated with ON/OFF state are plotted in phase space.

### 2.2 FIGR: Classification-based inference

In FIGR, we exploit the separability of the regulatory and kinetic parameters to break up the inference problem into two distinct tractable subproblems. For inferring the parameters of any given gene, we classify the data points into two classes—one in which the gene’s product is being synthesized (ON class) and the other in which the product is not being synthesized (OFF class). The regulatory parameters are inferred by determining the optimal *G* − 1 dimensional hyperplane separating the two classes. The kinetic parameters can be inferred either by fitting the piece-wise linear Glass equations to estimates of the rate of change of gene product concentrations or by fitting Equation 5 to the gene product concentration time series.

#### 2.2.1 Determining ON/OFF state

We will assume that the gene product concentration, including initial concentration, is bounded by the maximum concentration determined by the synthesis and degradation rates, that is,

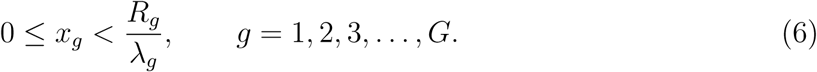

Let

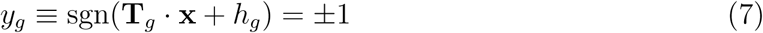

represent the ON/OFF state of gene *g*. Then,

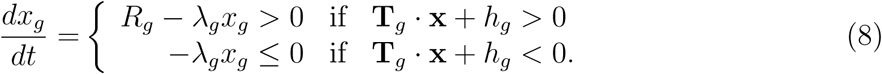

This implies that the ON/OFF state of a gene can be determined by ascertaining the sign of the *velocity*, 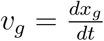.

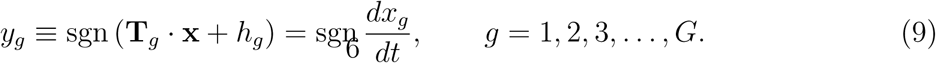

Gene expression data, such as those obtained from immunofluorescence or high-throughput sequencing, inevitably contain noise. If the gene expression level is close to its maximum 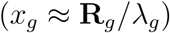 or minimum level 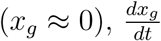 is theoretically close to zero, but noise causes sgn 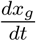 to fluctuate, which might be interpreted as spurious switching events. To avoid this problem, we identify a gene’s ON/OFF state as follows. If the gene expression level *x*_*g*_ is increasing (decreasing) at a rate greater than a user-supplied *velocity threshold* 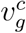, then the gene is classified as ON (OFF). Otherwise, if the expression level is above (below) a user-supplied *expression threshold* 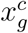, then the gene is classified as ON (OFF). This can be summarized as

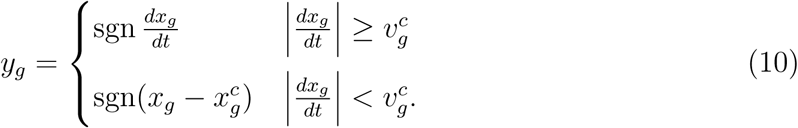

In our implementation of FIGR, cubic smoothing splines are fit to time series data and differentiated to estimate velocity. Figure 1E,H illustrates the determination of *y*_*g*_ for an example two-gene network.

#### 2.2.2 Determining regulatory parameters

Within the Glass model, the ON/OFF state of a particular target gene *g*, whose index we shall omit from now on, is given by *y* = sgn (**T** · **x** + *h*). Suppose that gene product concentrations have been sampled *P* times, in time and in one or more conditions or cell types. The gene ON/OFF state *y*_*p*_ is determined for each experimentally measured state vector **x**_*p*_, *p* = 1, 2, 3, …, *P*, according to the method described above (Section 2.2.1). Then, the regulatory parameters can be inferred by finding 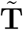 and 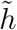 such that

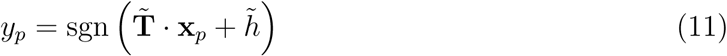

is satisfied for as many *p* as possible. Inferring the regulatory parameters therefore reduces to the problem of linear binary classification [14].

There are many well known supervised learning algorithms for linear binary classification. We have used both support vector machines (SVM) and logistic regression. An SVM finds a hyperplane buffered by the biggest possible margin such that the number of points **x**_*p*_ belonging to each class, “gene ON” or “gene OFF”, is maximized on opposite sides of the margin zone. This can be accomplished by minimizing the cost function

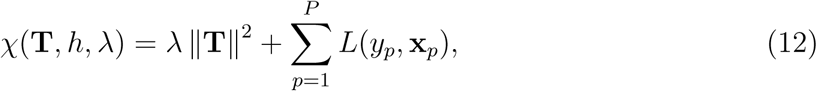

where the first term is a regularization penalty that maximizes the margin. The second term is the hinge loss function, *L*(*y*_*p*_, **x**_*p*_) = max (0, 1 − *y*_*p*_(**T** · **x**_*p*_ + *h*)), which is non-zero only for points that transgress their class boundary, each such point contributing an amount proportional to its distance from the margin. The parameter *λ* is used to choose the relative weight of the penalty and loss terms. Two-class logistic regression models the posterior probability of the ON/OFF state of a point as a logit transformation of its distance from the switching hyperplane. The optimal switching hyperplane can be found by minimizing Equation (12) with a Binomial deviance loss function 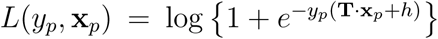. Figure 1C,F,I illustrate binary classification for an example two-gene network.

Minimization of Equation (12) is a convex optimization problem, which can be solved by quadratic programming or the Newton-Raphson method quite efficiently, even for large *G*. This is the key benefit of the separation of regulatory and kinetic parameters enabled by the Glass equations.

#### 2.2.3 Determining kinetic parameters

Having identified the ON/OFF state of a gene, *y*_*p*_, for *P* measurements of its concentration, *x*_*p*_, the Glass equations (Eq. 3) can be rewritten as

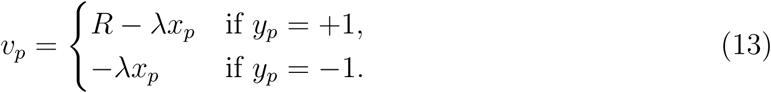

The velocities 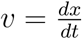 are estimated by differentiating cubic smoothing splines fit to the time series data (Section 2.2.1). Thus, for any particular gene, Equation (13) takes the form of *P* equations that are linear in the two unknowns *R* and *λ*. This is an overdetermined linear system, so best estimates for *R* and *λ* can be extracted by least-squares linear regression. In practice, the error in the spline, and hence in *v*, is the largest when a gene is switching states. We therefore exclude a user-configurable number of time points nearest to switching events. This method is implemented as the `“slope”` method of FIGR. Alternatively, *R* and *λ* can also be determined by fitting Equation 5 to the concentration data (see Section S1 of Supplementary Materials).

### 2.3 Inference of the *Drosophila* gap gene network

#### 2.3.1 Gene circuit equations for a spatially extended system

The gene circuit describes the time evolution of the concentrations of the gap proteins in a one-dimensional row of *N* nuclei lying along the anteroposterior axis of the embryo [24] during cleavage cycle 14. The lack of cell membranes in the syncitium implies that proteins can diffuse between nuclei (cells) and Equation (1) is modified to incorporate discretized Fickian diffusion and anteroposterior position, so that

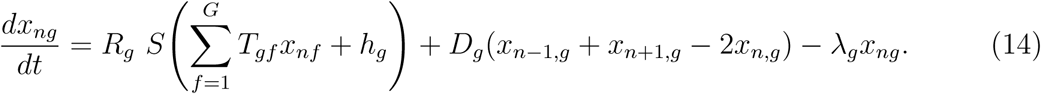

Here *x*_*ng*_(*t*) is the expression level of protein *g* in nucleus *n* at time *t, D*_*g*_ is the diffusion constant for protein *g*, and *S*(*u*) is the sigmoid regulation-expression function (Eq. 2). Zero-flux boundary conditions are used at the ends of the modeled region, while initial conditions are given by the empirical data from cycle 13.

#### 2.3.2 Model and spatiotemporal data

We inferred a gene circuit for the gap genes *hunchback* (*hb*), *Krüppel* (*Kr*), *giant* (*gt*), and *knirps* (*kni*). The model includes the upstream regulators Bicoid (Bcd), Caudal (Cad), and Tailless (Tll), so that *G* = 7. Other gap genes expressed in the head and tail of the embryo [15] are not included in the model, and so it is restricted to *N* = 58 nuclei lying between 35%–92% egg length (EL).

The gene circuit was inferred from a publicly available whole-mount immunofluorescence dataset [35] of the spatiotemporal expression of the segmentation genes. Data from the 50 minute-long cleavage cycle 14 are staged into 8 time classes, T1–T8, spaced 6.25 mins apart. We utilized integrated data [24] obtained by subtracting background fluorescence from raw single-embryo data, aligning the spatial patterns of different embryos belonging to a time class, and averaging over several embryos in each time class. See Janssens *et al*. [19] for details of how the data were processed.

#### 2.3.3 Classification-based Inference

The determination of ON/OFF state and inference of regulatory parameters **T**_*g*_ and *h*_*g*_ is carried out as described above in Sections 2.2.1 and 2.2.2. In addition to *R*_*g*_ and *λ*_*g*_, spatially extended gene circuits have an additional set of kinetic parameters, the diffusion constants *D*_*g*_.

We exploited so-called kink solutions to the gene circuit equations [36] to estimate the kinetic parameters. Let gene *g* be ON in an anteroposterior domain [*l, r*] so that there is net diffusion out of the domain into surrounding OFF nuclei. Then the balance of synthesis, degradation, and diffusion will establish a stable gradient,

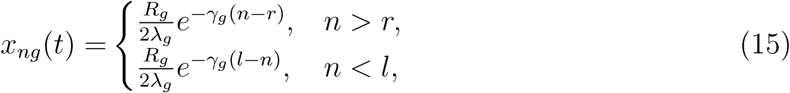

outside the domain. Here, 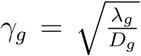 is the characteristic length scale of the gradient at steady state.

In order to infer the kinetic parameters, we first determine the time class in which a gene is expressed at the highest level, since the spatial gradient would best approximate steady state at that time point. Next, we identify gene-expression domains as local maxima in the spatial pattern in that time class. For genes having multiple expression domains, such as *hb* and *gt*, the domain with the highest expression is chosen. The gene is expressed at half maximum at the domain borders (Eq. 15). Therefore, border positions (*l* or *r*) are determined as nuclei where the expression is half of the domain peak. *R*_*g*_, *λ*_*g*_, and *D*_*g*_ are then determined by fitting Equation (15) to the observed expression data for *n > r* and *n < l* using MATLAB’s `lsqnonlin` function. This is implemented as the `“kink”` method of FIGR.

#### 2.3.4 Refinement of parameters for smooth gene circuits

For gene circuits with diffusion, the correspondence between velocity *v*_*g*_ and ON/OFF state *y*_*g*_ (Eq. 9) will be violated for a few nuclei lying adjacent to nuclei in which the gene is ON. Such nuclei would have positive velocity despite the gene being OFF because of diffusion of the protein from adjacent “ON nuclei”. Although this effect is limited to a few nuclei, it would nevertheless result in slightly inaccurate parameter estimates. For this reason, we refine the parameters inferred by FIGR using local optimization. The objective function is 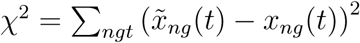, where 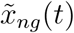 are data and *x*_*ng*_(*t*) are solutions of Equation (14). *χ*^2^ was minimized with MATLAB’s `fminsearch`, which utilizes the Nelder-Mead algorithm to conduct an unconstrained local search starting from parameters inferred with FIGR.

#### 2.3.5 Comparison with simulated annealing and timing

Simulated annealing was carried out with gene circuit C code as described previously [24]. Wall clock time was measured with C’s `time` function or MATLAB’s `tic`/`toc` functions for simulated annealing and FIGR respectively.

### 2.4 Validation of FIGR with synthetic data

Random gene circuits were generated and simulated to generate synthetic data as follows. *R*_*g*_ and *λ*_*g*_ were drawn uniformly from the interval [0.5, 2]. We chose *T*_*gf*_ and *h*_*g*_ such that the switching hyperplane passed through a random point **x**^cen^drawn uniformly from the bounding hypercube 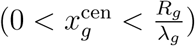, and the normal to the switching hyperplane 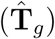 was drawn uniformly from the unit hypersphere. We generated *N* trajectories starting at random initial position **x**_*n*_(*t* = 0) drawn uniformly from the bounding hypercube by integrating the Glass equations without diffusion (Eq. 3) using MATLAB’s `ode45` solver. We stored the values of these functions *x*_*ng*_(*t*_*k*_) at 41 timepoints *t*_*k*_ = 0, 0.05, *…,* 2 to serve as synthetic data for FIGR.

## 3 Results

### 3.1 Validation of FIGR on synthetic data

As a first test of FIGR, we tested its ability to recover known parameters from synthetic data (see Section 2.4 for details). In each test, 10, 000 randomly generated gene circuits were simulated using the diffusion-less Glass equations (Eq. 3). For each gene circuit, *N* trajectories starting from random initial starting points were computed and sampled at 40 time points to obtain synthetic time series data. The parameters were inferred from the synthetic data using FIGR (Section 2.2) and compared with the known values by computing the discrepancies, 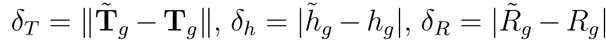, and 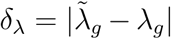, where 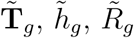, and 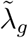 are inferred parameter values. The user-defined options and parameters utilized in this study are provided in Table S1 (Supplementary Material).

Figure 2A shows the frequency distributions of the discrepancies for 10, 000 two-gene networks. Given that **T**_*g*_ and 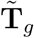 are unit normal vectors to the theoretical and inferred hyperplanes respectively, *δ*_*T*_ gives the angle between the two normals when it is small, while 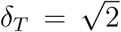 implies that the inferred hyperplane is orthogonal to the theoretical one. The switching hyperplane has been inferred accurately (*δ*_*T*_ < 0.1 or 6° angle to the theoretical normal) for the vast majority of random gene circuits. The inference of *h*_*g*_ and the kinetic parameters is similarly accurate. The inference of the genetic interconnectivity depends on the number of sampled data points and the number of modeled genes as one would expect (Fig. 2B). The accuracy improves with increasing number of trajectories (*N)* and decreases with increasing number of genes (*G*) if the number of data points is held constant.

**Figure 2:**
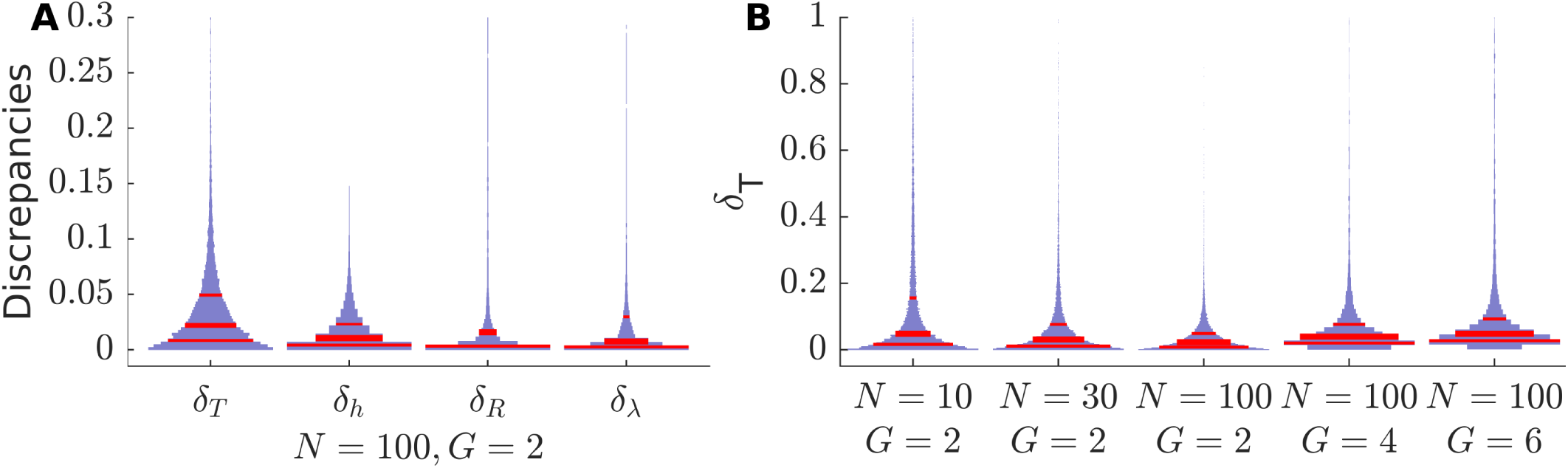
Tests of FIGR using synthetic data. Frequency distributions of the discrepancy between theoretical and inferred parameters for random gene circuits are shown as violin plots. The width of the violin plot is proportional to frequency and the red lines indicate first quartile, median, and third quartile. *G* is the number of genes in each circuit. *N* is the number of trajectories, starting from random points, that were simulated to generate synthetic data. **A**. Discrepancies for 10, 000 random two-gene gene circuits. **B**. Discrepancy in the normal to the switching hyperplane (*δ*_*T*_) as it depends on the number of genes and the number of sampled trajectories.

Although the inference is quite accurate, the discrepancies are not zero for most gene circuits and can be fairly large for a small number of gene circuits. This results from constraints imposed by the intrinsic dynamics of gene circuits and finite sample size. For instance, trajectories move away from the switching hyperplane for autoactivating genes. In this case, the initial conditions act as support vectors for the inferred hyperplane, which then strongly depends upon the random sample of starting points. Another situation that arises is that of a hyperplane that divides the bounding hypercube into ON and OFF regions in a lopsided manner. Since initial points are sampled uniformly, this results in too few sampled points in the vicinity of the hyperplane and poor inference. These considerations are also valid when inferring GRNs from empirical data. Such insights and their implications for parameter identifiability will be described elsewhere. Notwithstanding these constraints, our analysis demonstrates that FIGR is capable of inferring parameters quantitatively when provided with a sufficient number of data points.

### 3.2 Gap gene network inference

We tested the efficacy of FIGR on empirical data by inferring a gene circuit for the gap gene network acting during *Drosophila* segmentation. We provide a very brief summary of the segmentation system here; a more complete description may be found in reviews by Akam [1] or Jaeger [18]. The segmentation proteins pattern the anteroposterior axis during the first three hours of embryogenesis. During this period, the embryo is a syncitium, so that nuclei lack cell membranes and undergo 13 mitotic divisions, termed cleavages. After the tenth cleavage cycle, the majority of the nuclei migrate to the periphery of the embryo and are arranged in a monolayer, forming a syncitial blastoderm. Cellular membranes form by invagination of the plasma membrane in the latter half of cleavage cycle 14, at the end of which the embryo undergoes gastrulation.

Near the end of cleavage cycle 14, the segmentation genes are expressed in spatially resolved patterns that specify the position of each cell to an accuracy of one cell diameter. Segmentation gene expression is initiated by shallow protein gradients formed by the translation of localized mRNAs, such as *bicoid* (*bcd*) and *caudal* (*cad*), deposited in the oocyte by the mother. These maternal protein gradients regulate the gap genes, which commence mRNA expression in cleavage cycle 10–12 and are expressed in broad domains ∼20 nuclei wide during cycle 14. The gap genes in turn regulate the pair-rule and segment-polarity genes that provide the molecular prepattern for the subsequent segmentation of the embryo. All the maternal and gap proteins are known to act as transcription factors that regulate each others’ expression in a complex GRN that has been modeled extensively [3, 12, 13, 15– 17, 24, 25, 32, 36–38], providing an ideal test of FIGR.

We inferred a gene circuit for the gap genes *hunchback* (*hb*), *Krüppel* (*Kr*), *giant* (*gt*), and *knirps* (*kni*) regulated by the upstream factors *bcd, cad*, and *tailless* (*tll*). The equations utilize the sigmoid regulation-expression function and include the effects of the spatial diffusion of gap proteins (Section 2.3.1). The gene circuit models the protein expression of these genes between 35%–92% egg length during cleavage cycle 14 (Section 2.3.2). The dataset is comprised of immunofluorescence protein concentration measurements at the resolution of individual nuclei at eight timepoints during cleavage cycle 14 (Fig. 3A).

**Figure 3:**
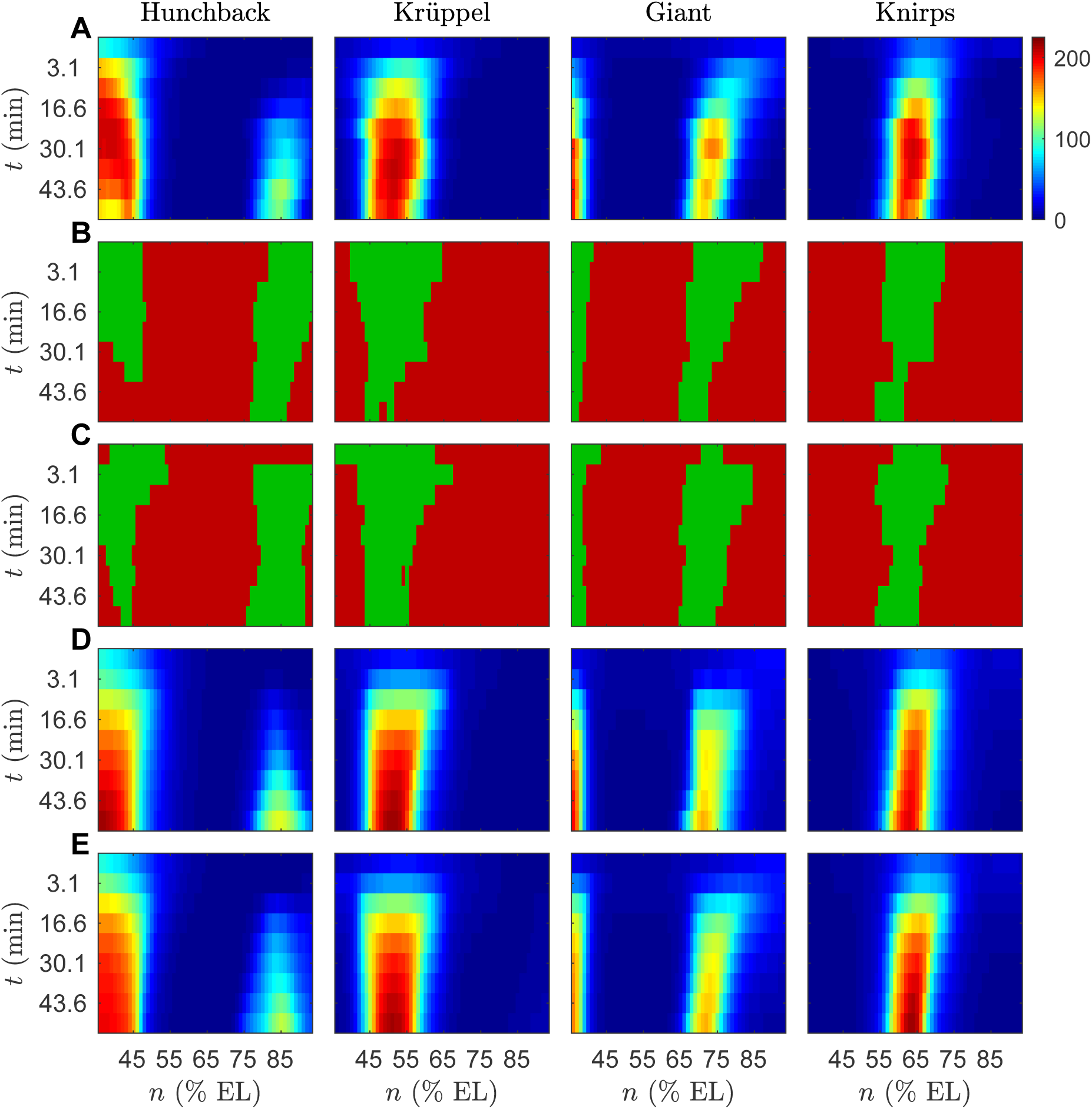
Classification-based inference of the *Drosophila* gap gene network. Horizontal axes indicate anteroposterior position. Vertical axes represent time elapsed since the thirteenth nuclear division in the downward direction. **A**. Integrated spatiotemporal gap gene expression data *x*_*ng*_(*t*_*k*_), where *k* = 1, *…,* 9, used to infer the gene circuit. Data at the first time point are from cycle 13 and serve as initial conditions for the model. The rest of the time points correspond to eight time classes in cycle 14 separated by 6.25 minutes. Hot (cold) colors indicate high (low) expression levels. **B**. Classification of ON/OFF state. Green (red) colors indicate ON (OFF) states *y*_*ng*_(*t*_*k*_) determined from velocities *v*_*ng*_(*t*_*k*_). **C**. Prediction of ON/OFF state 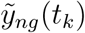 using parameter values inferred with FIGR. **D**. Gene circuit output using parameters obtained by FIGR and then refined using local search (RMS=13.29). **E**. Gene circuit output using parameters inferred by simulated annealing (RMS=11.03).

The regulatory and kinetic parameters were inferred with FIGR (Section 2.3.3). The user-defined options and parameters utilized for fitting the gap gene circuit are provided in Table S1 (Supplementary Material). Figure 3B shows that the assignment of ON/OFF states using slope estimates (Section 2.2.1) correctly recapitulates the extent and dynamics of the gene expression domains. The regulatory parameters 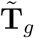 and 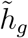 were inferred by supervised classification (Section 2.2.2). Predictions of the ON/OFF state from the inferred parameter values, 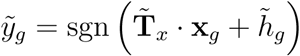 (Fig. 3C), are in good agreement with the empirical ON/OFF classification and the gene expression domains, suggesting correct inference of the regulatory parameters. In the presence of diffusion, the correspondence between velocity 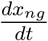 and the ON/OFF state of the gene will be violated for nuclei abutting gene expression domains due to diffusion of protein from ON regions into OFF regions. The inferred parameters likely deviate slightly from optimal values and were therefore refined using local search (Section 2.3.4). Gene circuit simulation with refined parameters produces output that agrees very well with the data and the output of gene circuits inferred using simulated annealing (Fig. 3D,E), and has an RMS 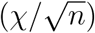 of 13.29.

In the inferred network (Fig. 4), the gap genes are activated by Bcd and Cad, repress each other, and autoactivate. The posterior gap genes are repressed by Tll. This scheme is in good agreement with experimental evidence [18], the network inferred by simulated annealing, and previous analyses [15, 24].

**Figure 4:**
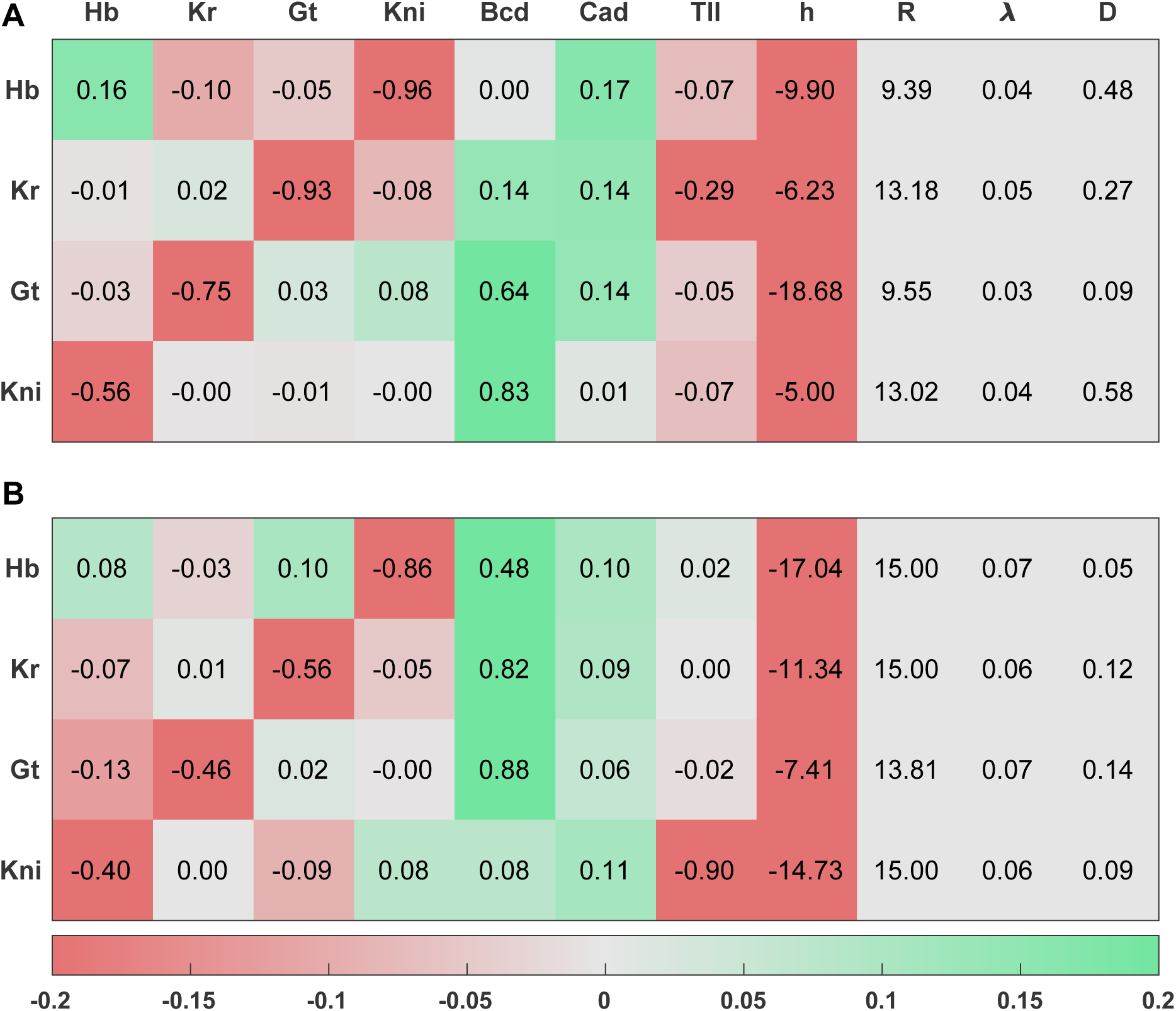
Comparison of gap gene parameters inferred by FIGR and simulated annealing. The values of 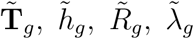, and 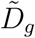 are listed one gene per row. The **T** matrix is shown in the first seven columns, one column per regulator. Green (red) indicates activation (repression). Since **T**_*g*_ and *h*_*g*_ can only be determined up to a constant factor, they have been normalized 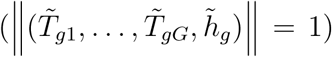 to allow comparison between FIGR and simulated annealing. **A**. FIGR after refinement. **B**. Simulated annealing.

With compiled MATLAB code, FIGR with refinement takes 48 sec compared to 29,067 sec taken by simulated annealing (∼600-fold speed up) to achieve the same RMS. In summary, FIGR infers the gap gene network accurately but at a fraction of the computational expense.

## 4 Discussion and Conclusions

Gene circuits [15, 24, 31] provide many unique advantages for inferring and modeling gene regulatory networks. Among higher-throughput statistical approaches for the inference of GRNs, some, such as ARACNe [26] and Module Networks [33], are limited to the inference, do not infer causality, and are incapable of predicting gene expression levels. Others, such as Dynamic Bayesian Networks [20], although capable of simulation and prediction, rely on the assumption that gene regulation by multiple TFs is additive and also cannot predict gene expression at intermediate time points. In contrast, gene circuit equations are biologically realistic in representing gene regulation as a nonlinear switch-like function of TF concentrations. Gene circuits not only infer the topology of the network but the directionality (causality) and sign of the edges. Most importantly, gene circuits are not limited to inference but are capable of accurately simulating and predicting gene expression patterns. Finally, the use of differential equations allows gene circuits to compute transient solutions, an important factor in simulating development since fate determination can occur before equilibrium is reached [25, 34].

Despite the promise held by gene circuits, their application, as of other nonlinear differential equation based models, has been limited to smaller networks so far. A major challenge in broader adoption is the high computational expense of inferring the free parameters from time series data. Currently, the approach for inferring parameter values [5, 31, 37] is to solve (“integrate”) the ODEs to obtain trajectories, compare with experimental trajectories, and refine parameters using global optimization techniques such as simulated annealing. This procedure is slow and expensive because it requires performing multidimensional optimization on a complicated objective function *χ*^2^(*{T, h, R, λ}*) with many local minima and each function evaluation involves solving a system of ODEs. For instance, inferring a relatively small 4-gene network takes 35 mins on 64 CPUs [37]. The computational complexity grows as *O*(*G*^3^), since *G* ODEs are solved in each function evaluation and the number of objective function evaluations required is proportional to the number of parameters (*O*(*G*^2^)).

In contrast, FIGR directly attempts to fit the differential equations, which describe how the velocities *v*_*g*_ depend upon the positions *x*_*g*_. **T**_*g*_ and *h*_*g*_ are determined using binary classification (support vector machines or logistic regression). Both of these algorithms reduce to quadratic programming, and thence to convex optimization. Subsequently, *R*_*g*_ and *λ*_*g*_ can be determined by linear regression against velocities or non-linear regression against concentrations using the piece-wise analytical solutions of the ODEs, which are even simpler optimization problems. Each inference can be completed in a fraction of a second on a consumer-grade computer, even with interpreted MATLAB code. We found that in the gap gene circuit, the values of the regulatory parameters **T**_*g*_ and *h*_*g*_ thus inferred were consistent with those obtained by simulated annealing, and ODEs solved with the inferred parameters produced gap gene domains correctly positioned in space and time. Furthermore, the parameter values, which were inferred under the assumption of a Heaviside regulation-expression function, served as a good starting point for equations with a sigmoid regulation-expression function. Gene circuits with a good RMS could be obtained with an “off-the-shelf” Nelder-Mead simplex method built into MATLAB. This refinement takes under a minute in serial compared to the eight hours taken by simulated annealing. Finally, logistic regression has a computational complexity of *O*(*G*), where *G* is the number of features (genes), which implies that FIGR has a complexity of *O*(*G*^2^) when inferring *G* genes. Consequently, FIGR scales with problem size much better than global nonlinear optimization techniques.

Besides the problem of computational efficiency, the broader application of gene circuits, and indeed all nonlinear differential equation models, is limited by a lack of understanding of parameter identifiability. Most commonly, *a posteriori* confidence intervals for the parameter estimates are computed [2]. Such calculations are based on the strong assumption that the solution is linear in the parameters and that the measurement errors are normally distributed. *a posteriori* parameter identifiability analysis also does not provide any hints to improve experimental design for achieving better identifiability in future studies.

Although we have not directly addressed the problem here, we anticipate that conceiving of and visualizing gene circuit inference as a classification problem will lead to insights into parameter identifiability. For example, it is evident from phase space plots (Fig. 1F,I) that sampling gene expression trajectories closer to the true switching hyperplane of the gene will lead to less “wiggle room” for the inferred hyperplane and result in more accurate parameter inference. This implies that datasets that measure gene expression near steady state, for example in differentiated cell types, are unlikely to lead to accurate parameter inference irrespective of the number of data points sampled or the precision of the experiment. Sampling transient trajectories densely when genes are turning ON or OFF is the best strategy for accurate parameter inference. Another less obvious implication is that it is easier to infer the regulation of negatively autoregulated genes than positively autoregulated ones. Trajectories move toward or away from the switching hyperplane for negatively or positively autoregulated genes respectively, making it more likely that sampled data points will lie near the hyperplane in the former. This analysis will be reported elsewhere.

In summary, we have exploited features of the mathematical structure of the Glass model to break a difficult optimization problem into a series of two, much simpler, optimization problems. We have demonstrated that FIGR is effective on synthetic as well as experimental data from a biologically realistic GRN. We have validated the inferred gap gene model by comparing its parameters against models inferred with simulated annealing as well as comparing its output against experimental data. The improvement in computational efficiency and scalability should allow the inference of much larger GRNs than was possible previously.

