## Supplementary Material for "Classification-based Inference of Dynamical Models of Gene Regulatory Networks"

David A. Fehr[1], Manu[2,*], and Yen Lee Loh[1,*]

[1]Department of Physics and Astrophysics,
[2]Department of Biology,
University of North Dakota,
Grand Forks, ND 58202, U.S.A.

[*]Equal contribution.

### S1 Alternative method for determining kinetic parameters

In the diffusion-less case, in addition to fitting the Glass equations to velocity data (Eq. 13), the kinetic parameters $R$ and $\lambda$ can also be determined by fitting Glass equation solutions (Eq. 5) to the concentration time series data. We identify time intervals during which all $y$ are either positive or negative so that

$$x_m(t_k) = \begin{cases} x_m(t_0)e^{-\lambda \Delta t_k} + \frac{R}{\lambda}(1 - e^{-\lambda \Delta t_k}) & \text{if } y_m(t_k) = +1 \quad \forall k, \\ x_m(t_0)e^{-\lambda \Delta t_k} & \text{if } y_m(t_k) = -1 \quad \forall k, \end{cases} \tag{S1}$$

where $m$ and $k$ index the time intervals and the time points lying inside a particular interval respectively. Within a particular interval, $x_m(t_k)$ is the concentration at the $k$th time point, $x_m(t_0)$ is the initial concentration, and $\Delta t_k = t_k - t_0$ is the time elapsed from the start of the interval. Equations S1 are $P \gg 2$ non-linear equations with two unknowns, $R$ and $\lambda$, and can be fit relatively easily using off-the-shelf non-linear optimization methods. We used MATLAB's `lsqnonlin` function that implements a Trust-Region algorithm. This is implemented as the "`conc`" method of CBI.

| Description | Option/Parameter | Acceptable Values | Values Utilized | |
|---|---|---|---|---|
| | | | Synthetic Data Tests | Gap Gene Inference |
| **Determining regulatory parameters** | | | | |
| Spline smoothing parameter for determining velocities | `splinesmoothing` | $[0, 1]$ | 1 | 0.01 |
| Velocity threshold for determining on/off state | `slopethresh` ($v_g^c$) | $\geq 0$ | 0.01 | 1 |
| Expression threshold for determining on/off state | `exprthresh` ($x_g^c$) | $> 0$ | 0.2 | 100 |
| **Determining kinetic parameters** | | | | |
| Method for determining the kinetic parameters | `Rld_method` | `'slope'` `'kink'` `'conc'` | `'slope'` | `'kink'` |
| **Determining kinetic parameters by "slope" method** | | | | |
| Margin to exclude unreliable velocity estimates near maxima and minima of the time series | `Rld_tsafety` | $\geq 0$ | 3 | NA |
| **Determining kinetic parameters by "kink" method** | | | | |
| Spline smoothing parameter for identifying spatial expression domains and border positions | `spatialsmoothing` | $[0, 1]$ | NA | 0.5 |
| Expression threshold above which points are included in fitting the kink equations, expressed as fraction of maximum domain expression | `minborder_expr_ratio` | $(0, 1)$ | NA | 0.01 |

Table S1: User-defined options and parameters utilized in FIGR code. The spline smoothing parameter is passed to the spline-fitting `csaps` function of MATLAB. It takes values between $0$ and $1$, where $1$ implies no smoothing while $0$ results in a straight-line fit.
